# Pulvinar influences parietal delay activity and information transmission between dorsal and ventral visual cortex in macaques

**DOI:** 10.1101/405381

**Authors:** Yuri B. Saalmann, Ryan Ly, Mark A. Pinsk, Sabine Kastner

**Affiliations:** Department of Psychology, University of Wisconsin-Madison, 1202 West Johnson Street, Madison WI 53706; Princeton Neuroscience Institute, Princeton University, Washington Road, Princeton NJ 08544; Department of Psychology, Princeton University, Washington Road, Princeton NJ 08544

**Author notes:** Contributed equally (Yuri B. Saalmann, Ryan Ly). Corresponding Authors* Yuri B. Saalmann, Department of Psychology, University of Wisconsin-Madison, 1202 West Johnson Street, Madison WI 53706, Ryan Ly, Princeton Neuroscience Institute, Princeton University, Washington Road, Princeton NJ 08544.

**Keywords:** Thalamus, lateral intraparietal area, V4, attention, feedback, synchrony, oscillation

## Abstract

The fronto-parietal attention network represents attentional priorities and provides feedback about these priorities to sensory cortical areas. Sustained spiking activity in the posterior parietal cortex (PPC) carries such prioritized information, but how this activity is sustained in the absence of feedforward sensory information, and how it is transmitted to the ventral visual cortical pathway, is unclear. We hypothesized that the higher-order thalamic nucleus, the pulvinar, which is connected with both the PPC and ventral visual cortical pathway, influences information transmission within and between these cortical regions. To test this, we simultaneously recorded from the pulvinar, lateral intraparietal area (LIP) and visual cortical area V4 in macaques performing a selective attention task. Here we show that LIP influenced V4 during the delay period of the attention task, and that the pulvinar regulated LIP-V4 information exchange. Pulvino-cortical effects were consistent with the pulvinar supporting sustained activity in LIP. Taken together, these results suggest that pulvinar regulation of cortical functional connectivity generalizes to dorsal and ventral visual cortical pathways. Further, the pulvinar’s role in sustaining parietal delay activity during selective attention implicates the pulvinar in other cognitive processes supported by such delay activity, including decision-making, categorization and oculomotor functions.

**Significance Statement:** A network of areas on the brain’s surface, in frontal and parietal cortex, allocate attention to behaviorally relevant information around us. Such areas in parietal cortex show sustained activity during maintained attention and transmit behaviorally relevant information to visual cortical areas to enhance sensory processing of attended objects. How this activity is sustained and how it is transmitted to visual areas supporting object perception is unclear. We show that a subcortical area, the pulvinar in the thalamus, helps sustain activity in the cortex and regulates the information transmitted between the fronto-parietal attention network and visual cortex. This suggests that the thalamus, classically considered as a simple relay for sensory information, contributes to higher-level cognitive functions.

## INTRODUCTION

Higher-order thalamic nuclei, like the pulvinar, form prevalent cortico-thalamo-cortical pathways, in contrast with first-order thalamic nuclei, like the lateral geniculate, which form major pathways from the sensory periphery to the cortex (1-5). The pulvinar has been shown to regulate neural activity across the ventral visual cortical pathway in macaques, i.e., areas V1, V4, TEO and TE (6-8). This includes modulating the gain of cortical neurons (6) and neural synchrony within and between cortical areas (7, 8) during the delay period of an attention task or stimulus-evoked responses. In contrast, little is known about the pulvinar’s influence on neural activity in the dorsal visual cortical pathway through the posterior parietal cortex (PPC).

In macaque PPC, the lateral intraparietal area (LIP) is involved in representing attentional priorities, oculomotor processing, decision-making and categorization (9-12). LIP neurons show stimulus-evoked responses as well as characteristic delay period activity, i.e., increased spike rate during maintained attention, action planning or decision-making (in the absence of visual stimulation) relative to baseline (9, 10). While feedforward inputs to LIP can account for early stimulus-evoked responses, it is not clear how LIP activity is sustained during delay periods.

Because the pulvinar has reciprocal connections with the dorsal pathway (13-15), it is well positioned to influence LIP activity. We hypothesized that the pulvinar influences delay period activity of LIP neurons (hypothesis 1) based on three findings. First, studies in rodents suggest that higher-order thalamic nuclei, the mediodorsal nucleus and the motor thalamus, are crucial for sustained neuronal firing in frontal cortex (16-18). Second, in macaques, there is strong pulvinar influence on the cortex during delay periods along the ventral visual cortical pathway (7). Finally, deactivation of the dorsal pulvinar, which is interconnected with LIP, produces severe deficits in visually-guided actions (19), similar to damage to the PPC.

LIP also has been shown to provide feedback to dorsal extrastriate cortex in order to modify the gain of MT neurons (20, 21), which is important for selective attention and other cognitive functions. LIP is also connected to the ventral visual cortical pathway (22, 23). However, the relationship between neural activity in LIP and ventral extrastriate cortex has not been directly tested. We hypothesized that LIP influences V4 during spatial attention (hypothesis 2). In light of the fact that the frontal cortex can influence V4 (24, 25), an additional parietal influence on V4 would allow for greater flexibility in the attentional modulation of V4 responses, e.g., LIP providing information on stimulus salience and the frontal eye fields (FEF) providing internally-generated goal-directed information (26).

A further question is whether the pulvinar regulates activity between the dorsal and ventral visual cortical pathways, as available anatomical evidence suggests that the lateral pulvinar connects with both the PPC and higher-order areas of the ventral visual cortical pathway, e.g., V4, TEO, and TE. The PPC tends to have more connections dorsally, and the ventral visual cortex more ventrally, in the lateral pulvinar as well as pulvinar as a whole (5, 27, 28). Because the lateral pulvinar has been shown to regulate interactions within the ventral visual cortical pathway, we hypothesized that the lateral pulvinar also regulates interactions between dorsal and ventral visual cortical pathways (hypothesis 3).

To test our hypotheses, we simultaneously recorded from interconnected sites in the pulvinar, LIP and V4 while macaques performed a selective attention task. A number of pieces of evidence support pulvinar influences on LIP delay activity (hypothesis 1): the pulvinar contained a subset of cells with shorter response latencies than the bulk of LIP cells; and attentional modulation of pulvinar spike-LIP field coherence correlated with LIP delay activity. Further evidence supports interactions between LIP and V4 (hypothesis 2) as well as the pulvinar regulating the information transmission between LIP and V4 (hypothesis 3): the pulvinar influenced both LIP and V4 in overlapping frequency ranges, in which LIP and V4 also interacted, based on spike-field coherence and Granger causality estimates.

## RESULTS

We report simultaneous recordings of single-unit spiking activity and local field potentials (LFPs) from three areas: the pulvinar (n=51 cells and 56 LFPs), LIP (n=41 cells and 56 LFPs) and V4 (n=31 cells and 56 LFPs), in two macaques performing a flanker task. This task allowed us to manipulate the monkey’s spatial attention, because a spatial cue appeared randomly at one of six different locations, drawing the monkey’s attention to the upcoming target position in a circular array of barrel and bowtie shapes (both monkeys >80% correct performance overall; Fig. 1). For each monkey, we used anatomical connectivity maps derived from diffusion MRI, as well as overlapping receptive fields (RFs) for recording sites, to guide electrode placements in interconnected thalamo-cortical networks.

**Fig. 1.**
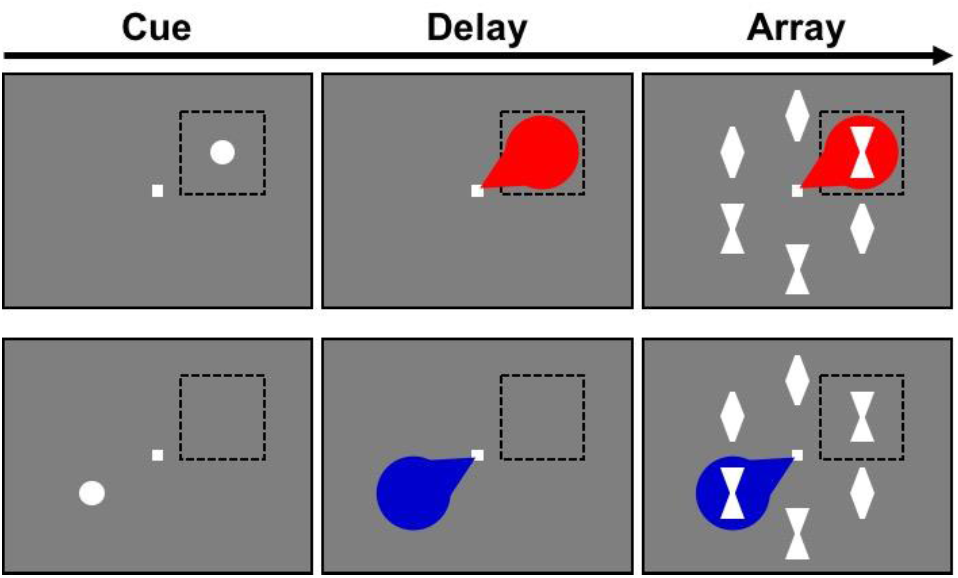
Flanker task used to manipulate spatial attention. Spatial cue (white filled circle) signals location of target in forthcoming stimulus array. Target can be positioned inside (top right) or outside (bottom right) the RF (black square). The monkey’s attention (red/blue spotlight) is necessarily drawn to, and maintained at, the cued location, for correct identification of the target (here, a bowtie).

### Response latencies in LIP, V4 and pulvinar

The dorsal pathway (LIP) needs to interact with the ventral pathway (V4), to integrate spatial attention and visual object information. As a first step towards characterizing information flow between the dorsal and ventral pathways, we calculated the response latency in LIP, V4, as well as in the pulvinar, which is connected with both areas (Fig. S1). The median latency of the neuronal population in each area in response to the cue was 51 ms (n=36), 51 ms (n=29) and 65 ms (n=46) for LIP, V4 and the pulvinar, respectively (consistent with previous reports of response latency in LIP, e.g., (29), V4, e.g., (30, 31) and pulvinar, e.g., (32)). In response to the target embedded in the array, the median onset latency was 50.5 ms (n=36), 46.5 ms (n=30), and 56.5 ms (n=42), respectively. Although cortical areas showed shorter median latencies than the pulvinar, a subset of pulvinar neurons – 30% for cue-evoked responses and 43% for array-evoked responses – responded earlier than half the neurons in LIP and V4 (consistent with hypothesis 1; Fig. 2). Taken together, this suggests that a subset of cortical neurons initially provides visual and/or attentional information to a subset of pulvinar neurons (also see SI Appendix, Fig. S2). Next, this subset of pulvinar neurons helps recruit additional cortical neurons into the activated neuronal ensemble representing the behaviorally relevant information. Based on the long latencies (>100ms) of subsets of cortical and pulvinar neurons, such pulvino-cortical interactions may repeat.

**Fig. 2.**
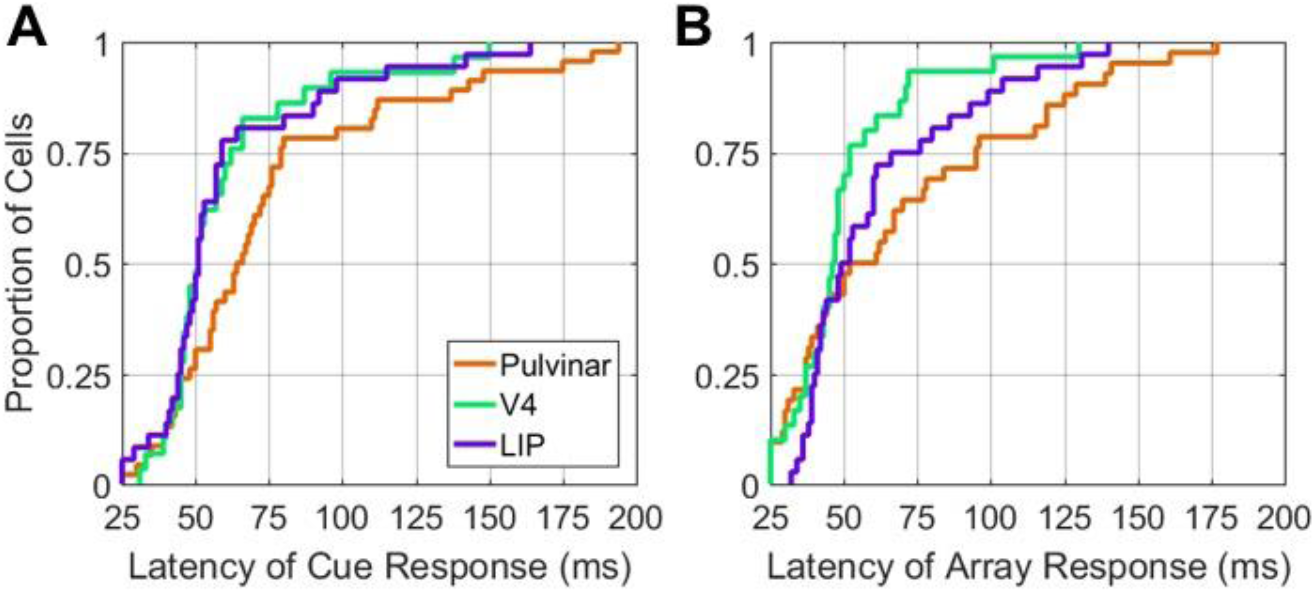
Comparison of response latencies between the pulvinar, V4, and LIP. Cumulative distribution function of response latencies for the pulvinar, V4, and LIP in response to the cue at the RF **(A)** and target at the RF within the array **(B)**.

### Delay period spiking activity in LIP, V4 and pulvinar

Delay period spiking activity is a signature of many neurons in LIP (9, 10, 12) and has also been reported in other areas, including V4 (33-35) and pulvinar (7). There was significantly increased delay period activity (0-200ms before array onset) observed in LIP neurons (n=41; p=0.00034, Wilcoxon signed-rank test; Fig. 3A), pulvinar neurons (n=51; p=0.0046; Fig. 3B) and V4 neurons (n=31; p=0.00002; Fig. 3C) at the population level, when attention was directed at neuronal RFs (as compared to when attention was directed away from the RF). For an individual neuron, there did not appear to be any clear preference for spiking at one particular time during the delay period of individual trials; rather, there was variable spike timing from trial-to-trial across the delay. To probe the underlying mechanisms contributing to this attention-enhanced delay period activity in LIP, V4 and pulvinar, in the following sections we measured cortico-cortical and thalamo-cortical interactions across the delay period.

**Fig. 3.**
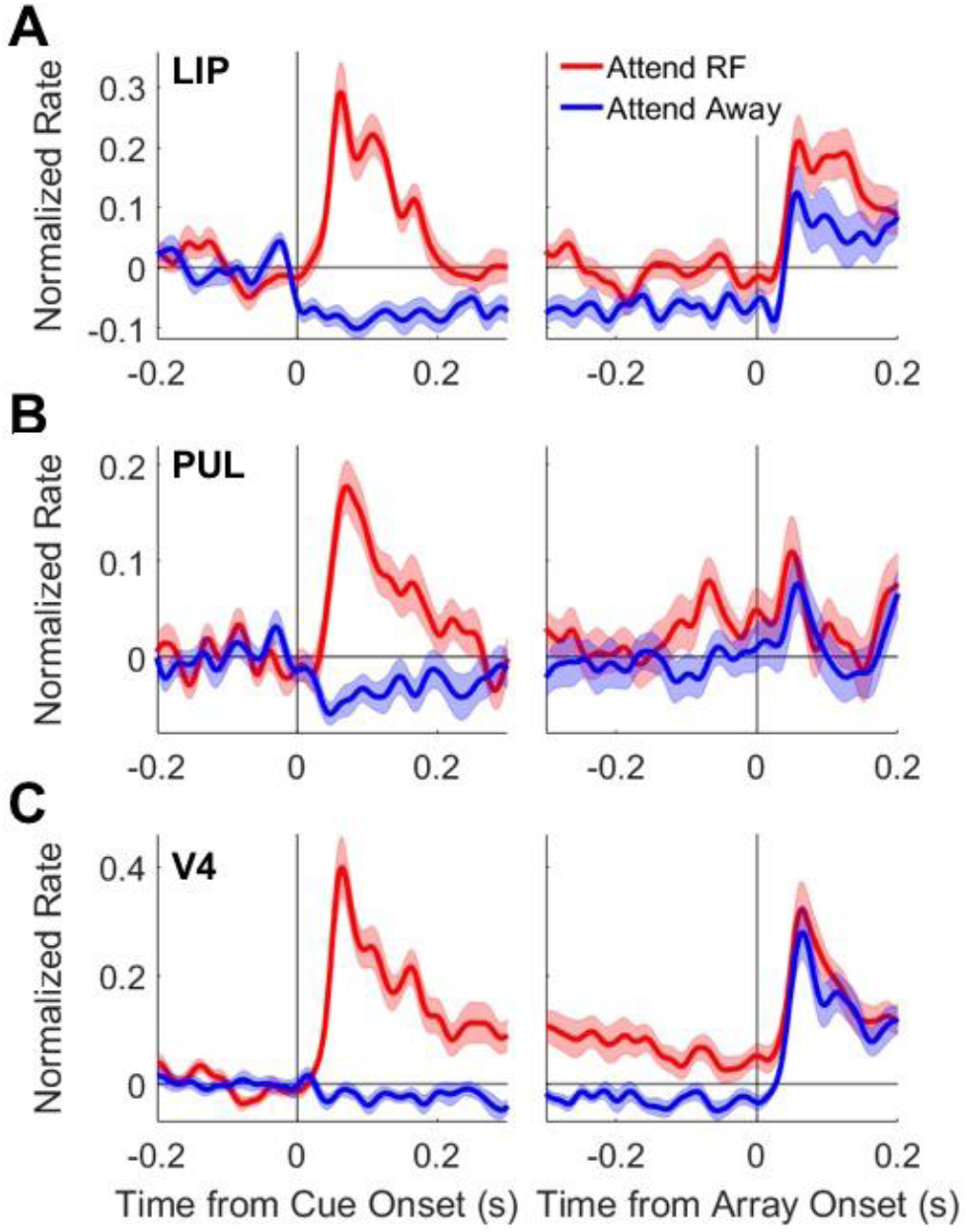
Delay period spiking activity in pulvinar, V4 and LIP. Population spike density functions for LIP (**A**), pulvinar (**B**), and V4 (**C**). LIP, pulvinar and V4 neurons show increased spike rate in the delay period between cue onset and array onset when attention was directed at the RF (red) compared to attention away from the RF (blue). All error bars are SEM.

### LIP influence on V4

The putative salience map in LIP (10) is well positioned to provide attentional feedback to V4. Spatial attention increased LIP influence on V4 during the delay period, but attention did not significantly change V4 influence on LIP. At the population level, there was significantly increased LIP spike-V4 field coherence with attention (versus attention away) in the alpha (8-15 Hz; n=38; p=0.015; Wilcoxon signed-rank test), beta (15-30 Hz; p=0.006) and gamma frequency ranges (30-50 Hz; p=0.0005; Fig. 4 A and D). This shows a linear dependency between LIP output (spikes) and V4 input (reflected in the LFP). We next calculated a statistical measure of causality between LIP and V4. Conditional Granger causal influence of LIP on V4 (accounting for pulvinar influence) also significantly increased with attention in the gamma range (n=56 p=0.013, t test; alpha range, p=0.096; beta range, p=0.081; Fig. 4 C and F). In comparison, there was no significant increase in conditional Granger causal influence of V4 on LIP or significant increase in V4 spike-LIP field coherence in these frequency ranges (Fig. S3). Considering the delay period activity observed in V4 (Fig. 3C), these data suggest that the dorsal visual cortical pathway provides information about attentional priorities to the ventral visual cortical pathway, to selectively modulate V4 neuronal excitability (supporting hypothesis 2). Spatial attention also increased within-LIP spike-field coherence in the gamma range (n=40, p=0.002; Fig. 4 B and E), consistent with attention adjusting the degree of synchrony between LIP cells. This suggests that LIP feedback to V4 can be modulated by adjusting LIP spike rate or LIP neural synchrony.

**Fig. 4.**
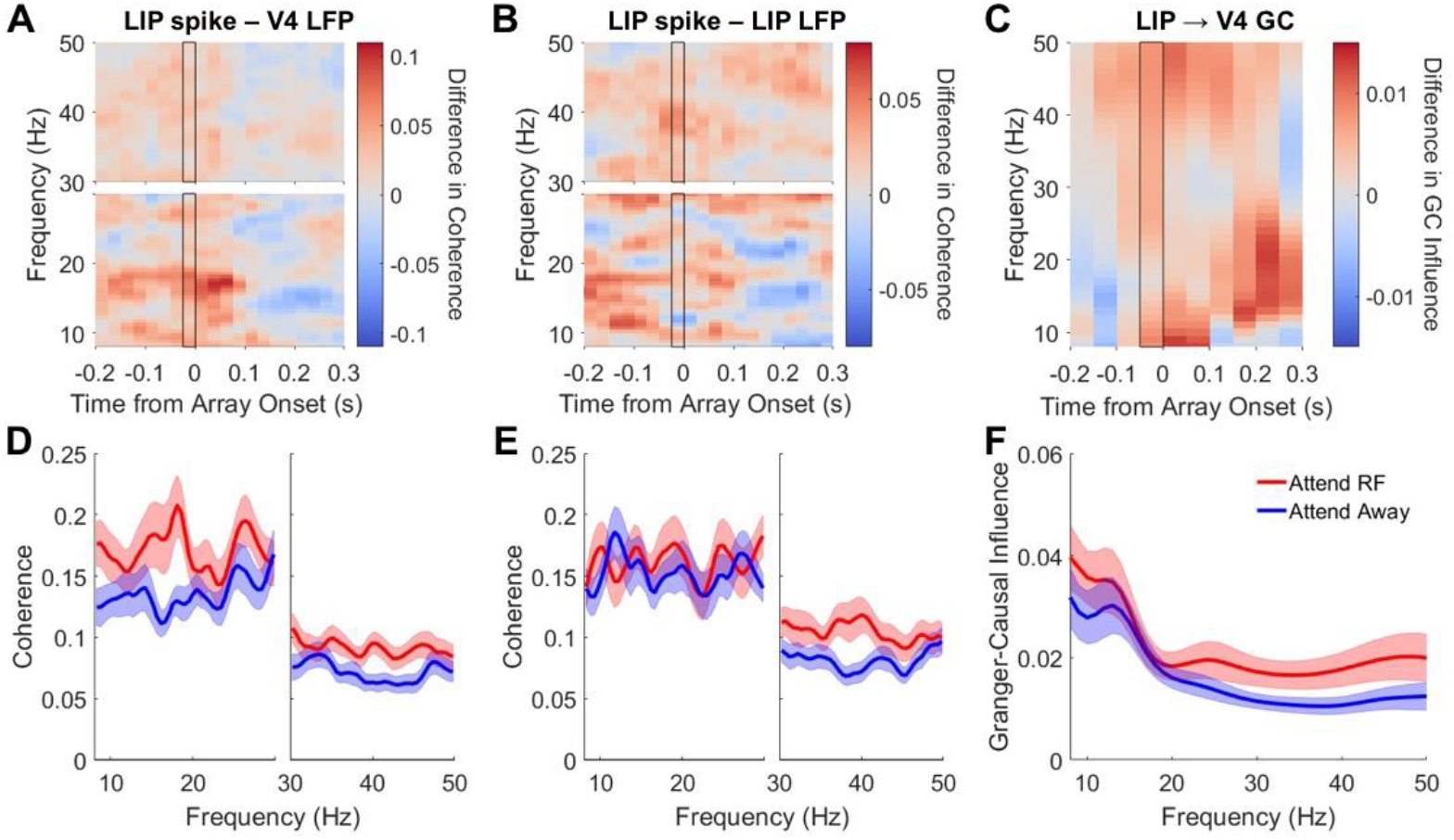
LIP influence on V4. **(A-C)** Time-frequency plots of **(A)** LIP spike-V4 field coherence, **(B)** within-LIP spike-field coherence, and **(C)** conditional Granger causal influence of LIP LFP on V4 LFP (accounting for pulvinar). Plots represent attention at RF condition minus attention away from RF condition. Spectra calculated in 300 ms sliding windows, in 25 ms steps. Data are aligned to array onset. **(D-F)** Spectra calculated in 0- 300 ms window prior to array onset (black-outlined column in A-C). All error bars are SEM. (**D**) LIP spike-V4 field coherence. (**E**) Within-LIP spike-field coherence. (**F**) Conditional Granger-causal influence of LIP LFP on V4 LFP (accounting for pulvinar).

### Pulvinar influence on information transmission between LIP and V4

The pulvinar has previously been shown to influence information transmission along the ventral visual cortical pathway, including V4 (7, 8). If the pulvinar regulates information transmission between the dorsal and ventral visual cortical pathways, then this might be done simply by the pulvinar influencing both LIP and V4 in an overlapping frequency range. At the population level, there was significant pulvinar spike-LIP field coherence in the alpha (n=32, p=0.005, Wilcoxon signed-rank test) and beta (p=0.0009) ranges (Fig. 5 A and D), as well as significant pulvinar spike-V4 field coherence in the beta range (n=36, p=0.006), and trending in the alpha range (p=0.054; Fig. 6 A and C), with attention at neuronal RFs (versus attention away) during the delay period. Similarly, spatial attention increased conditional Granger causal influence of the pulvinar on LIP (accounting for V4) in the alpha (n=56, p=0.00039, t test) and beta (p=0.0063) ranges (Fig. 5 C and F; consistent with hypothesis 1), as well as the pulvinar on V4 (accounting for LIP) in the alpha (n=56, p=0.007) and beta (p=0.0117) ranges (Fig. 6 B and D). The pulvinar influenced both LIP and V4 in the alpha and beta ranges, which would allow for LIP-V4 communication through coherence at these frequencies (supporting hypothesis 3). This is consistent with the aforementioned LIP spike-V4 field coherence in the alpha and beta ranges. Within-pulvinar spike-field coherence also increased in the beta range (n=44, p=0.013, Wilcoxon signed-rank test; alpha range, p=0.08; Fig. 5 B and E) with attention, suggesting that the pulvinar influenced the cortex by increasing the synchrony as well as spike rate of pulvinar neurons.

**Fig. 5.**
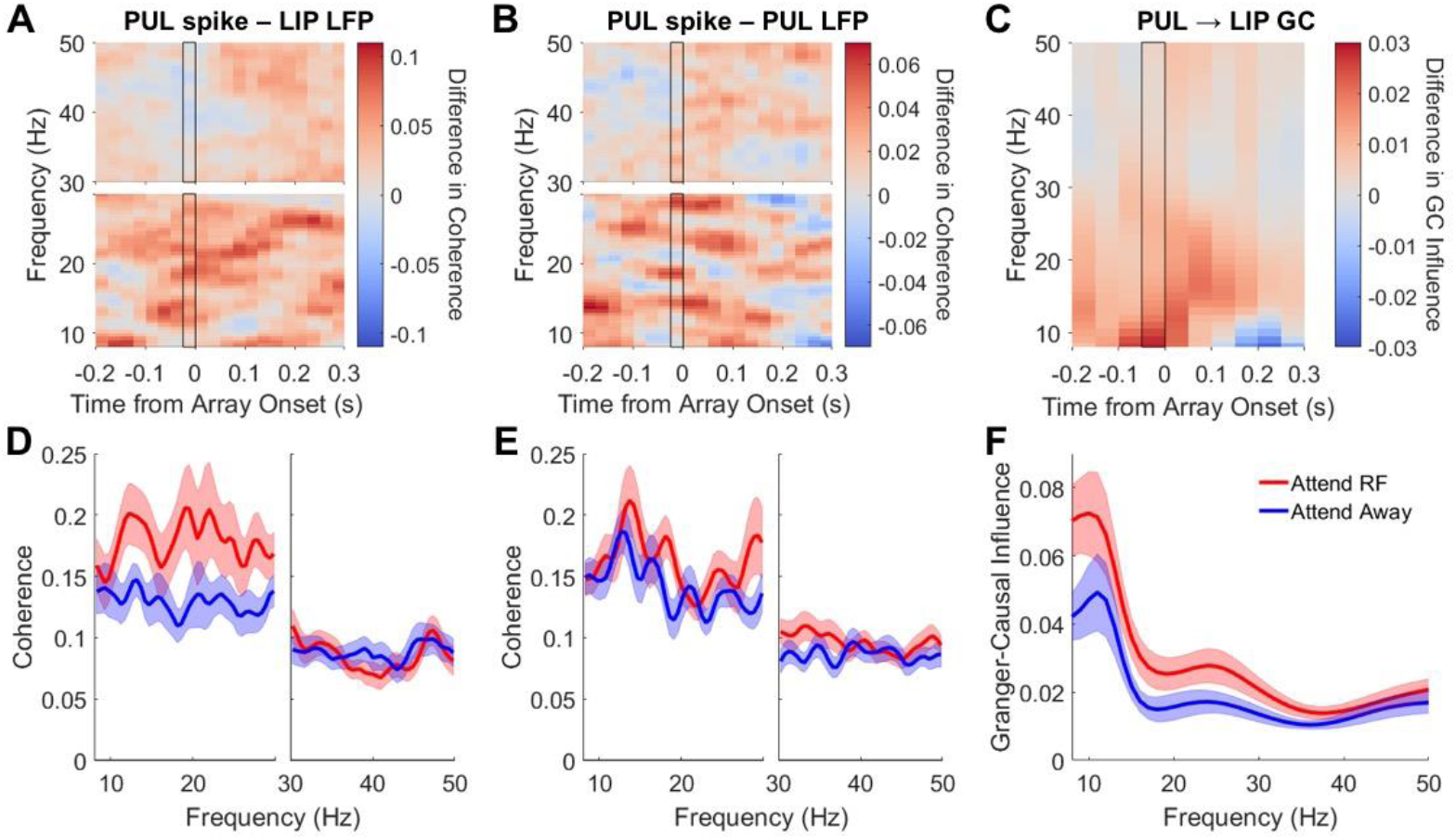
Pulvinar influence on LIP. **(A-C)** Time-frequency plots of **(A)** Pulvinar spike-LIP field coherence, **(B)** within-pulvinar spike-field coherence, and **(C)** conditional Granger causal influence of pulvinar LFP on LIP LFP (accounting for V4). Plots are as in Fig. 4. **(D-F)** Spectra calculated in 0-300 ms window prior to array onset (black-outlined column in A-C). (**D**) Pulvinar spike-LIP field coherence. (**E**) Within-pulvinar spike-field coherence. (**F**) Conditional Granger-causal influence of pulvinar LFP on LIP LFP (accounting for V4).

**Fig. 6.**
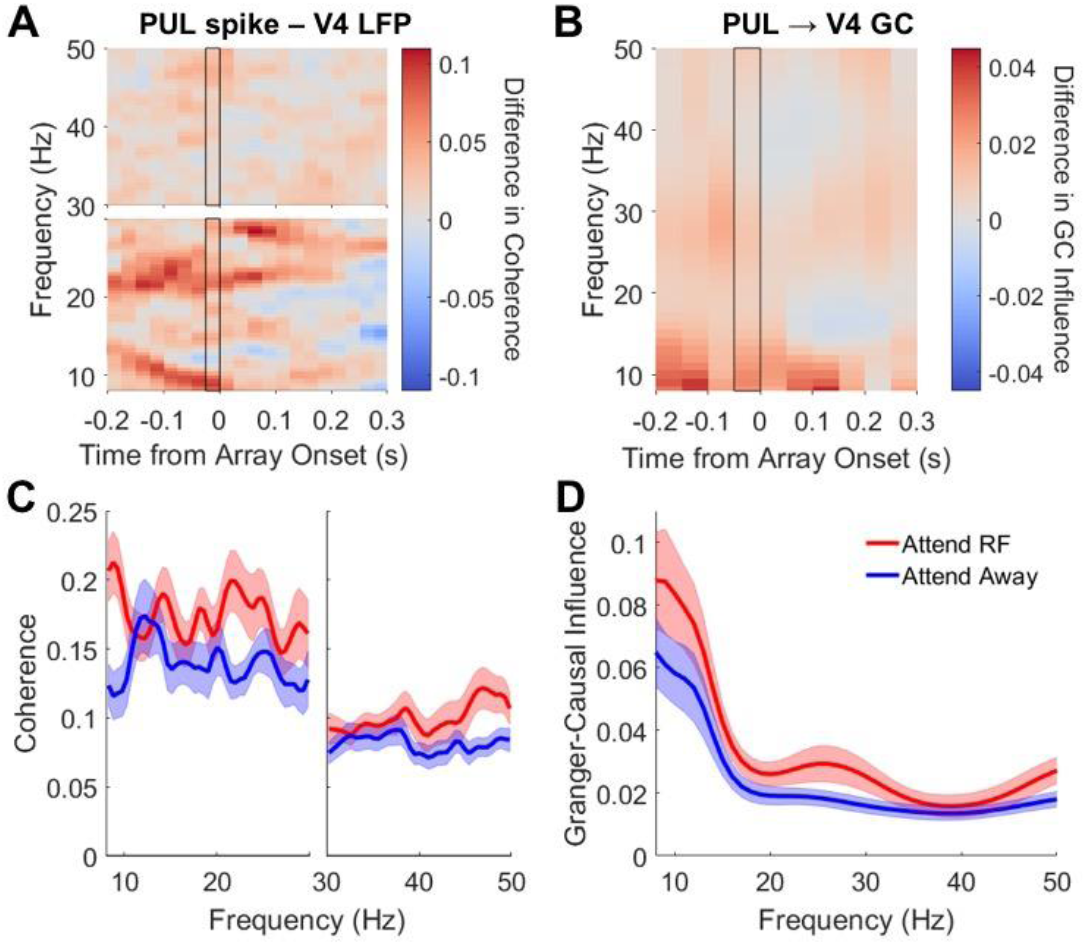
Pulvinar influence on V4. **(A-B)** Time-frequency plots of **(A)** Pulvinar spike-V4 field coherence, and **(B)** conditional Granger-causal influence of pulvinar LFP on V4 LFP (accounting for LIP). Plots are as in Fig. 4. **(C-D)** Spectra calculated in 0-300 ms window prior to array onset (black-outlined column in A-B). (**C**) Pulvinar spike-V4 field coherence. (**D**) Conditional Granger-causal influence of pulvinar LFP on V4 LFP (accounting for LIP).

Tracer studies suggest that the dorsal pulvinar predominantly connects with the dorsal cortical pathway, whereas the ventral pulvinar connects with the ventral visual cortical pathway (5, 28). This prompts the question of what zone(s) of the pulvinar, if any, regulates communication between dorsal and ventral cortical areas? Although tracer studies of pulvinar connections with LIP and V4 have not been performed in the same animal to the best of our knowledge, different studies show that both LIP and V4 connect with the lateral pulvinar (5, 13-15, 36). Considering the brachium of the superior colliculus as the demarcation between dorsal and ventral pulvinar, our probabilistic tractography on diffusion MRI data showed that dorsal pulvinar predominantly connected with LIP, whereas ventral pulvinar predominantly connected with V4, consistent with tracer studies. However, LIP and V4 projection zones did overlap in the intermediate region between dorsal and ventral pulvinar, particularly in lateral pulvinar (Figure 7). In comparison, the projection zones of two ventral visual cortical areas, TEO and V4, showed greater overlap in ventral pulvinar (5) (including in the same animals (7)). This suggests that an intermediate region between dorsal and ventral pulvinar mediates interactions between dorsal and ventral visual cortical areas, which would likely offer reduced wiring costs within the pulvinar.

**Fig. 7.**
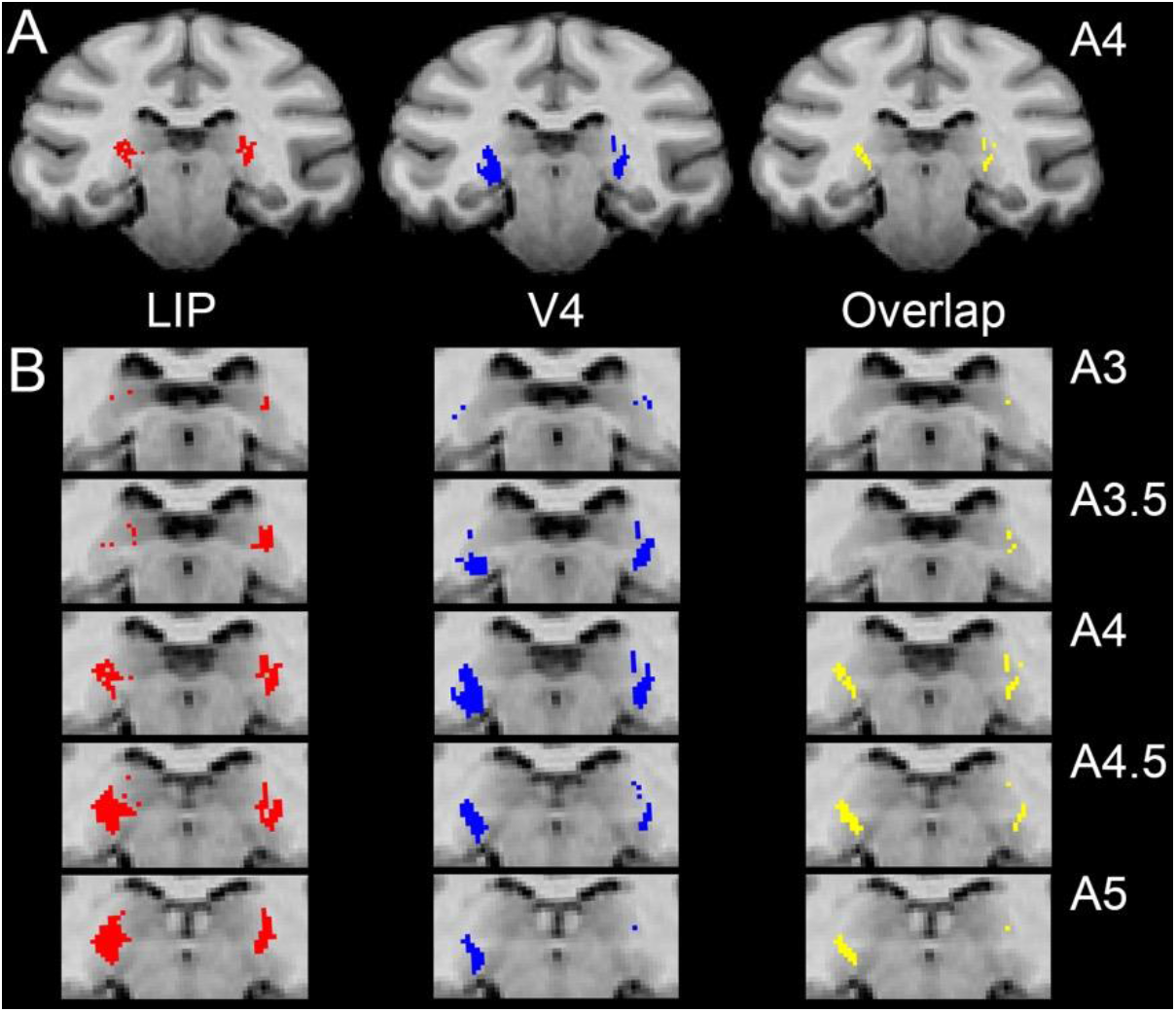
Overlapping projection zones of LIP and V4 in the pulvinar. (**A**) Pulvinar voxels connected with LIP (red), V4 (blue) or both (yellow) are shown overlaid on T1-weighted coronal slice at 4 mm anterior to interaural line. (**B**) Sequential slices (0.5 mm separation) zoomed in on pulvinar.

### Pulvinar influence on cortical delay period spiking activity

If the pulvinar influences LIP delay period activity, then one might expect coherence between pulvinar output (spikes) and LIP input (reflected in the LFP) to correlate with LIP spike rate during the delay period. Indeed, there was a significant positive correlation between attention-enhanced (i.e., difference between attention at RF and attention away) pulvinar spike-LIP field coherence in the alpha range and LIP spike rate during the delay period (0-300ms before array onset) (n=26; Spearman r=0.44, p=0.012; Fig. 8A). This suggests that the pulvinar supports the maintenance of LIP delay period activity (supporting hypothesis 1).

**Fig. 8.**
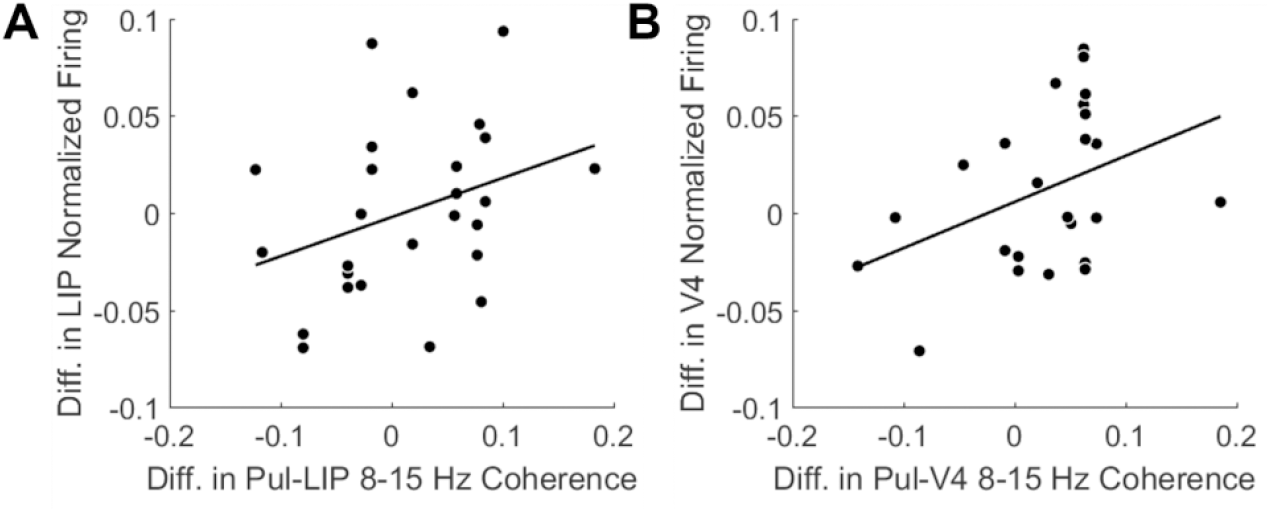
Pulvinar contributes to cortical delay activity. (**A-B**) Scatter plots showing relationship between attention-related change (difference between attention at RF and attention away) in coherence and spike rate. Coherence and spike rate calculated in 0- 300 ms window prior to array onset. Each point corresponds to data from an individual recording session. (**A**) Attention-related increase in pulvinar spike – LIP field alpha coherence correlates with increase in LIP delay firing rate. (**B**) Attention-related increase in pulvinar spike – V4 field alpha coherence correlates with increase in V4 delay firing rate.

Given that the pulvinar influenced LIP delay period activity, it prompts the question of whether the pulvinar influenced V4 delay period activity as well? There was a trending positive correlation between attention-enhanced pulvinar spike-V4 field coherence in the alpha range and V4 spike rate during the delay period (0-300ms before array onset) (n=24; Spearman r=0.38, p=0.033 but not significant after correcting for multiple comparisons; Fig. 8B). This suggests that pulvinar influence on cortical delay period activity may be a mechanism operating in both parietal and visual cortex.

## DISCUSSION

### Pulvinar contributions to cortical delay period activity (hypothesis 1)

Spatial attention increased pulvinar Granger causal influence on LIP and V4 in the alpha and beta frequency ranges. Attention also increased coherence between pulvinar spike output and cortical LFPs (in LIP and V4), and this pulvino-cortical interaction in the alpha range correlated with cortical delay period spiking activity. These results suggest that the pulvinar played a role in shaping and maintaining delay period spiking and LFP activity in parietal and visual cortex. There were at least three possible underlying mechanisms increasing the effect of pulvinar output on cortical excitability during spatial attention: first, pulvinar spike rate increased; second, pulvinar neurons synchronized, allowing for summation of post-synaptic responses in the cortex; and third, the pulvinar and LIP/V4 synchronized (within the 8-30 Hz range), increasing the likelihood that pulvinar spikes arrived in the cortex during periods of reduced inhibition.

Previous studies that pharmacologically manipulated the pulvinar in primates have shown that the pulvinar can strongly augment visual cortical activity during visual stimulation. In anesthetized prosimians (galago), stimulating the lateral pulvinar (with bicuculline) increased stimulus-evoked spiking activity of V1 neurons, when both the pulvinar and V1 RFs overlapped; and deactivating the pulvinar (with muscimol) had the opposite effect (6). In behaving macaques, deactivating the ventro-lateral pulvinar (with muscimol) reduced both visually-evoked V4 spiking activity and later attention-enhanced V4 activity during visual stimulation (8). Further evidence has also supported thalamic influence over cortical activity during the delay period in attentional and working memory tasks. Our previous macaque work, using the same flanker task as in this study, has shown that the pulvinar influenced LFPs (at frequencies >8 Hz) in the ventral visual cortical pathway (V4 and TEO) during maintained attention across the delay period in the absence of visual stimulation (7). Recent mouse studies have shown that other higher-order thalamic areas, specifically the mediodorsal thalamus and motor thalamus, contribute to the sustained spiking activity of frontal cortical neurons (16-18). Beta synchrony between the mediodorsal thalamus and frontal cortex was important for maintaining information across the delay (18). We extend these findings to show that the pulvinar contributes to the sustained activity of cortical neurons in primates. The relative response latencies between LIP, V4 and pulvinar neurons, as well as the attention-mediated increases in spike-field coherence (within the 8-30 Hz range) between pulvinar and cortex in both directions, suggest that a subset of cortical neurons may first activate a subset of pulvinar neurons, which in turn can either help activate additional cortical neurons, and so on, similar to the mechanism proposed for interactions between frontal cortex and mediodorsal thalamus (16, 18) (although the preferential spiking of an individual neuron at a particular time during the delay in the mouse studies, giving rise to a “tiling” of responses from different neurons across the delay period, was not clear in our study, possibly a species difference) or reciprocally excite the same cortical neurons. Such cortico-thalamo-cortical cycles could continue across the delay period until the appearance of a new visual stimulus.

LIP is part of the fronto-parietal attention network in macaques, and previous attention studies have shown that the response of LIP neurons to a visual stimulus and across a delay reflects attentional priorities (37), as in our task. Accordingly, PPC lesions in humans (38) and macaques (39) can give rise to severe attention deficits, such as perturbed attentional orienting to contralesional space, including visuo-spatial hemineglect. Because our study shows that the pulvinar contributes to LIP delay period activity, and likely PPC excitability more broadly, considering the anatomical connectivity between the pulvinar and PPC (5, 28), it should not be surprising that thalamic lesions involving the pulvinar in humans (40-42) and macaques (19, 43) can also produce deficits in directing attention to contralesional space. In addition to maintaining a representation of attentional priorities, PPC delay period spiking and LFP activity has been proposed to reflect working memory, accumulation of sensory evidence for decision-making and action planning (9, 12, 44). The pulvinar is well positioned to engage in all these cognitive functions, via its influence on PPC delay period activity.

### Interactions between dorsal and ventral visual cortical pathways (hypothesis 2)

LIP and V4 had similar median response latencies and latency distributions to the spatial cue in the flanker task and, during the following delay period, spatial attention only modulated LIP influence on V4. These results suggest that LIP and V4 both represented visual stimulus information at a similar time after stimulus onset, with LIP subsequently providing information on spatial attention priorities to V4. In a recent study where monkeys were cued to categorize stimuli based on either their motion or color (45), early (bottom-up) representation of visual information about the cue in both V4 and LIP (with no significant latency difference) was followed shortly after by a transient representation of the task-relevant information in V4, and later by a sustained (top-down) representation of the task-relevant information first in LIP and then in V4. These temporal dynamics were similar to that in our study, except our study suggests that LIP may extract salient task information without early V4 input, depending on task requirements (i.e., especially if largely spatial).

Spatial attention modulates V4 spiking activity (46-48) and neural synchrony, e.g., increasing gamma activity (49, 50). Previous work on attentional feedback to V4 has focused on frontal cortical sources (24, 25), e.g., electrically stimulating FEF modulates V4 spiking, and attention increases FEF synchrony with V4. However, lesioning the entire lateral prefrontal cortex, including FEF, in one hemisphere reduced, but did not eliminate, attentional modulation of ipsilateral V4 activity; and monkeys showed little decrement in behavioral performance (51). Our study shows that the PPC, particularly LIP, is another source of top-down attentional influence on V4. The LIP influence on V4 operated in the same frequency range, i.e. gamma, as FEF influence on V4 (24). However, we also found significant LIP spike-V4 field coherence at alpha and beta frequencies. Different frequency bands have been proposed to predominantly contribute to different functions, e.g., (52, 53), and processing in different cortical layers (54-56) (but see (57)). Longer-distance feedback pathways tend to target superficial layers and shorter-distance feedback pathways tend to target deep layers (58, 59). As LIP feedback pathways to V4 might be considered of intermediate distance (targeting both superficial and deep layers), it is possible that LIP gamma-frequency influence represents feedback to superficial layers in V4, and LIP-V4 alpha/beta interaction represents feedback to deep layers in V4 (60). This suggests a possible difference in the contributions of FEF and LIP to V4 processing, i.e., LIP may have greater influence over activity in deep layers of V4.

### Pulvinar regulation of LIP and V4 interactions (hypothesis 3)

The pulvinar influenced LIP and V4 activity at both alpha and beta frequencies, and LIP spikes synchronized with V4 LFPs at these frequencies. This is consistent with the pulvinar adjusting functional connectivity between LIP and V4. Our previous work has shown that pulvinar influence on LIP and V4 extends to the gamma range, via alpha-gamma cross-frequency coupling (7, 61). This suggests that the pulvinar may also regulate the LIP influence on V4 at gamma frequencies reported here. Such pulvinar influence over gamma-band cortico-cortical connectivity is supported by a recent macaque study showing that pulvinar deactivation (with muscimol) reduces gamma as well as beta coherence between LFPs in V4 and inferior temporal cortex (8). Taken together, these results indicate an important role for the pulvinar in shaping the pattern of rhythmic activity across a range of frequencies within and between LIP and V4, thereby enabling coordinated processing and information transfer between both areas.

It has been proposed that directly connected cortical areas are indirectly connected via the pulvinar – called the “replication principle” (5). Although there are direct connections between LIP and V4 (22, 23), evidence for indirect connections via the pulvinar is lacking. Our diffusion MRI data suggest that pulvino-LIP connections and pulvino-V4 connections partially overlap at an intermediate depth in the (lateral) pulvinar. Although diffusion MRI data does not have sufficient spatial resolution to identify individual neurons, the pulvinar control of LIP-V4 functional connectivity is consistent with a cortico-thalamo-cortical path carrying information from one cortical area to the pulvinar, in order to coordinate activity with the second cortical area.

The dorsal and ventral visual cortical pathways are commonly considered to preferentially represent spatial/intentional and object-based information, respectively. Previous attentional work has focused on pulvinar interactions with the ventral visual cortical pathway (7, 8). This study shows that the pulvinar not only interacts with the dorsal cortical pathway, but does so in a similar mechanistic manner with the ventral cortical pathway. Current data support the pulvinar increasing the gain of cortical neurons and regulating cortico-cortical coherence across a range of frequencies, to increase the efficacy of cortical transmission of behaviorally relevant information. The pulvinar may also reduce the efficacy of cortical transmission of irrelevant information through inhibitory mechanisms, as evidenced by its modulation of cortical alpha oscillations (53, 62). The pulvinar may thus amplify relevant and filter out irrelevant information in the world around us.

## MATERIALS AND METHODS

The Princeton University Animal Care and Use Committee approved all procedures, which conformed with the National Institutes of Health guidelines for the humane care and use of laboratory animals. The LIP data have not previously been published, but an analysis of much of the pulvinar and V4 data is in print (7). However, here we focus on new analyses and previously unpublished findings. We simultaneously recorded from LIP, V4 and pulvinar of two male macaques (*Macaca fascicularis,* 4-8 years old) performing a flanker task variant (7). Full methods can be found in Supplementary Information, Materials and Methods.

## ACKNOWLEDGEMENTS

This work supported through NIMH grant R01MH110311 (Y.S.), NIMH grant R01MH064043 (S.K.), NEI grant R01EY017699 (S.K.), NIMH Conte Center grant 1P50MH109429, and the James S. McDonnell Foundation 21st Century Science Initiative – Understanding Human Cognition – collaborative grant.

## Author contributions

Y.S. performed electrophysiology; Y.S. and M.P. performed neuroimaging; R.L. and Y.S. analyzed data; Y.S., R.L. and S.K wrote the paper.

## SUPPLEMENTARY INFORMATION

## RESULTS

### Cortical influence on the pulvinar

For the pulvinar to influence cortical excitability and the efficacy of information transfer according to task demands, it needs to integrate attentional and visual information from other areas. At the population level, both LIP spike-pulvinar field coherence (n=40, p=0.003, Wilcoxon signed-rank test; Fig. S2 A and D) and V4 spike-pulvinar field coherence (n=29, p=0.01; Figure 7B) significantly increased during the delay period when attention was at neuronal RFs (compared with attention away) in the beta frequency range. In light of the shorter median response latencies of LIP and V4 neurons relative to pulvinar neurons, these data are consistent with the pulvinar receiving visuo-spatial attentional information from both LIP and V4.

## MATERIALS AND METHODS

### Behavioral task

We trained two male monkeys (*Macaca fascicularis,* 4-8 years old) to perform a flanker task variant (1). Monkeys initiated trials by depressing a response lever after an auditory “go” signal. This triggered the appearance of a 0.5 ° square fixation point at the center of the monitor (eye-monitor distance = 57 cm). After a variable delay of 300-700 ms, a 1.5° circular spatial cue randomly appeared for 100 ms duration, at one of six possible stimulus locations. After another variable delay period of 400-800 ms, six barrel- or bowtie-shaped stimuli, each 4×2°, appeared equally-spaced in a circular array around the fixation point, for 700 ms duration or until the monkey released the response lever. We positioned the circular array such that at least one stimulus appeared in the receptive field (RF) of recorded neurons. On half the trials, the stimulus at the pre-cued location, the target, was congruent with its nearest neighboring stimuli, the distracters; i.e., each of these three stimuli was barrel-shaped, or each was bowtie-shaped. On the other half of trials, the target and its nearest distracters were incongruent; i.e., a barrel target was flanked by bowtie distracters, or vice versa. If the target was barrel-shaped, then the monkey needed to release the lever immediately for juice reward (150-650 ms after target onset). Conversely, if the target was bowtie-shaped, then the monkey needed to release the lever after the stimulus array disappeared (150-650 ms after array disappearance). Because the stimulus array contained equal numbers of barrels and bowties, the expected performance accuracy for random responses was 50%. In fact, the monkeys performed the task with greater than 80% accuracy overall, suggesting that they maintained attention at the cued location during the delay period until target presentation. To ensure that the monkey maintained fixation throughout trials, 10% of all trials were ‘catch’ trials, in which the fixation point disappeared at a random time, requiring the monkey to immediately release the lever. Trials aborted if the monkey broke fixation, i.e., if eye position deviated by more than one degree from fixation.

We controlled stimuli, response monitoring and rewards using Presentation software. We presented visual stimuli at 50% contrast (light gray on darker gray background) on a 21 inch cathode ray tube monitor set at a 100 Hz refresh rate. A customized photodiode system affixed to a second monitor receiving identical input enabled verification of visual stimulus timing. Monkeys manipulated a lever with their hands to report decisions and received juice reward via a tube connected to an infusion pump. We monitored eye position using an infrared camera, operating at 120 Hz, with an ASL eye-tracking system.

### Acquisition of Structural and Diffusion-Weighted Images

We anesthetized monkeys with Telazol (tiletamine/zolazepam, 10 mg/kg i.m.) and atropine (0.08mg/kg i.m.) during scan sessions. We positioned monkeys in a customized MRI-compatible stereotaxic apparatus, and monitored their respiration rate and pulse rate respectively using an MRI-compatible respiratory belt and pulse oximeter. We acquired images at a 3 T head-dedicated scanner using a 12-cm transmit-receive surface coil. Prior to the head implant surgery, we acquired diffusion-weighted images (DWI) using an eddy-current compensated double spin-echo, echo-planar pulse sequence (2-4), with 1.0 mm^2^ in-plane resolution and 60 different isotropic diffusion directions (5) (field of view (FOV) = 128 x 96 mm; FOV phase = 75%; matrix = 128 x 96; phase partial fourier = 6/8; no. of slices = 47; slice thickness = 1.1 mm; repetition time (TR) = 10,000 ms; echo time (TE) = 145 ms; *b*-values = 0 and 1,000 s/mm^2^; slice orientation = transverse; 12:1 ratio of DWI to non-DWI) (5, 6). Data acquisition included twenty 60-direction sets of diffusion-weighted data for subsequent averaging, matching in-plane gradient echo field map and magnitude images to perform geometric unwarping of the diffusion-weighted data (TR = 500 ms, TE = 6.53/8.99 ms, flip angle = 55°), and T1-weighted structural images for co-registration (Magnetization-Prepared RApid Gradient-Echo (MPRAGE); FOV = 128 mm^2^; matrix = 256 x 256; no. of slices = 128; slice thickness = 0.5 mm; TR = 2,500 ms; TE = 4.38 ms; flip angle = 8°; inversion time (TI) = 1,100 ms; in-plane resolution = 0.5 mm^2^). In a separate scan session, we acquired 12 T1-weighted structural images and calculated the average image for each monkey, to generate a higher-quality structural brain image.

### Electrophysiology

We surgically implanted a customized plastic recording chamber, affixed to the skull with titanium screws and self-curing acrylic, in monkeys anesthetized with isoflurane (induction 2-4%, maintenance 0.5-2%). Four 2.5 mm craniotomies drilled within the recording chamber provided access to our pulvino-cortical regions of interest (ROIs) in the right hemisphere. We fitted each craniotomy with a conical plastic guide tube filled with bone wax (1, 7), through which glass-coated platinum-iridium electrodes traversed. These guide tubes held electrodes in place between recording sessions. During recordings, we stabilized the animal’s head using four thin rods that slid into hollows in the side of the acrylic implant. We micropositioned electrodes in each ROI with electrode microdrives coupled to an adapter system, attached to the top of the recording chamber, allowing different approach angles for each ROI. We amplified and filtered (150-8,000 Hz for spikes; 3-300 Hz for LFPs) electrode signals (40,000 Hz sample rate for spikes; 1,000 Hz sample rate for local field potentials (LFPs)) using a preamplifier with a high input impedance headstage and Plexon Multichannel Acquisition Processor controlled by RASPUTIN software. Control recordings for LFP quality in each ROI with three different reference electrodes – either a skull screw, silver wire in contact with the dura, or electrode in the white matter just outside the ROI – yielded similar LFPs, so we used a skull screw as the reference electrode during recording sessions. We sorted spikes online to map the RF of isolated neurons, then re-sorted spikes offline using Plexon Offline Sorter software. We first plotted a neuron’s RF using hand-held stimuli, then confirmed the RF by systematically flashing visual stimuli around the RF location while the monkey fixated centrally. The reported cells and LFPs in each recording session had overlapping RFs.

### Probabilistic tractography on diffusion MRI data

We used FSL software to analyze diffusion MRI data (8, 9). We corrected DWI and non-DWI for eddy currents using affine registration (12 degrees of freedom (DOF), FMRIB’s Linear Registration Tool (FLIRT)) to a non-DWI reference volume, and averaged to improve the signal-to-noise ratio (10). Next, we geometrically unwarped images using field map and magnitude images acquired in the same session (11). That is, the magnitude image was skull-stripped using FMRIB’s Brain Extraction Tool (BET) (12), forward-warped using FMRIB’s Utility for Geometrically Unwarping EPIs (FUGUE), and registered (6 DOF) to an averaged, skull-stripped non-DWI reference volume. We applied the resulting transformation matrix to the field map image (scaled to rad/s and regularized by a 2-mm 3D Gaussian kernel), which was subsequently used to unwarp DWI and non-DWI with the FUGUE utility. We then skull-stripped the T1-weighted structural brain image and co-registered to the averaged, skull-stripped and geometrically unwarped non-DWI reference volume (12 DOF), to produce the transformation matrix between the two spaces.

For probabilistic diffusion tractography (PDT) analyses, we manually delineated LIP, V4 and pulvinar ROIs for the right and left hemisphere of each monkey. We used the individual monkey’s T1-weighted structural brain image, in conjunction with a stereotaxic atlas (13), to guide the definition of the ROIs. We applied the transformation matrix, derived from the co-registration of the structural image to the reference non-DWI, to the ROI masks for PDT analyses.

We performed tractography analyses using FMRIB’s Diffusion Toolkit (FDT). The tractography algorithm modeled two fiber populations per voxel (14), suited to the complex fiber architecture of the thalamus (1, 15). For each monkey, we calculated probability distributions of fiber direction at each voxel (15, 16). To identify pulvinar voxels with a high probability of connection with V4 and LIP, we performed a PDT analysis to estimate pathways passing through any voxel in a pulvinar seed, and the probability such pathways will pass through a voxel in either of the two cortical targets, V4 and LIP (i.e., FDT’s “single mask seed with classification targets” tractography). From each pulvinar seed voxel, 5000 samples were drawn from the probability distribution (0.2 curvature threshold, 0.25 mm step length), and the proportion of these samples passing through each cortical target equated to the probability of connection to that target. We applied a threshold removing voxels with a less than 5% of maximum connection probability with the target, then calculated the overlap between thresholded pulvinar volumes respectively connected to V4 and LIP.

### Imaging electrodes in situ

To verify electrode locations in the pulvinar, V4 (prelunate gyrus) and LIP, we acquired T1-weighted structural brain images with platinum-iridium electrodes held *in situ* by the customized guide tubes. Although the electrode itself is not visible in the T1-weighted images, a susceptibility “shadow” artifact appears along the length of the electrode with a width of approximately one voxel (0.5 mm^3^, either side of the electrode). Our experimental approach was to position electrodes at the most dorsal point of an ROI (for a particular dorsal-ventral trajectory), then acquire structural brain images. During subsequent recording sessions, we used a microdrive to lower electrodes through ROIs to isolate neurons and logged all recording site coordinates from the microdrive system. At the end of an electrode track, i.e., at the most ventral point of our ROI, we acquired additional structural brain images, before starting a new track. We reconstructed the position of the electrode for each recording session, using the structural images of the start and end of each track as well as the daily microdrive coordinates.

### Spike rate analysis

We calculated spike density functions, convolving each spike with a 10 ms Gaussian and averaging across trials. Next, we subtracted baseline activity (200 ms before cue onset) from the response for each condition. Finally, to normalize responses, we divided the response by the maximum firing rate of any condition. For statistical analysis of delay period activity, we computed the mean across the 200 ms period before array onset. To contrast attention to the neuronal RF versus attention away, the neuronal RF is defined as the cue location that evoked the largest firing response 25-200 ms after cue onset; the attention away location is defined as the cue location that evoked the smallest firing response 25-200 ms after cue onset.

### Spike latency analysis

We detected the first peak or trough greater than 2 standard deviations from baseline (200ms before cue onset), in the 25-200 ms period after cue and array onset. We calculated spike response latency as the time to half-peak. Cells with responses that did not meet the criteria were excluded. For computing firing rate peaks in response to the array, if activity in the delay period (200 ms before array onset) was less than the baseline activity, the delay activity was used in place of the baseline. For computing firing rate troughs in response to the array, if activity in the delay period was greater than the baseline activity, the delay activity was used in place of the baseline.

### Spike-field coherence analysis

We used the coherence measure to study the temporal relationship between all possible spike-LFP combinations involving LIP, V4 and pulvinar. The coherency is given by 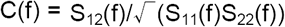, where S(f) is the spectrum with subscripts 1 and 2 referring to the simultaneously recorded spike train and LFP. For each paired spike-LFP recording, we calculated the spike-field coherence in 300 ms sliding windows (25 ms steps across the trial) for each attention condition: attention at the RF location or attention away from the RF. Attention at the RF condition reflected the RF of the spiking neuron; however, there was overlap between this RF and the LFP response field. The random location of stimuli from trial-to-trial result in attention conditions from any one recording session having an unequal number of trials. Because the number of trials affects the coherence estimate, we bias-corrected/transformed coherence values (17). The transformed spike-field coherence, T(f), is given by T(f) = tanh^-1^(C(f))-1/(V_0_-2), where V_0_ is the degrees of freedom. For our multi-taper estimates, v_0_ = 2*K*N, where K is the number of tapers (3) and N is the number of trials. To obtain population values, we averaged the transformed coherence estimates. To control for spikes affecting the LFP, we excised 2 ms around each spike time from the raw data trace and linearly interpolated these segments of the data trace. Because the results of LFP analyses were the same regardless of whether spikes were excised or not (in the frequency range of interest), we reported LFP data without spike excision. For all spectral analyses, we mainly focused on the delay period after the evoked response until the array onset, because during this period the monkey maintained spatial attention and the data in each session generally satisfied methodological assumptions of stationarity (18). To compare attention conditions across the population, we used Wilcoxon signed-rank tests to determine whether there was significantly greater coherence in particular frequency bands (i.e., alpha, beta and gamma) when attention was at the RF location compared to when attention was away from the RF. For this and all other spike/LFP analyses, we controlled the experiment-wise error rate (p < 0.05) using the Holm’s sequential Bonferroni procedure, and reported p values that survived this correction for multiple comparisons, unless otherwise specified.

To measure coherence in different frequency bands with sufficient frequency resolution, we first bandpass filtered data into alpha (8-15 Hz), beta (15-30 Hz), and gamma (30-50 Hz) bands using FIR filters (Kaiser window; model order 3960, transition bandwidth 1 Hz, stopband attenuation 60 dB, passband ripple 0.01 dB). We calculated spike-field coherence in the 300 ms period before array onset, using the Chronux toolbox for Matlab (http://chronux.org/ (19). For frequencies greater than 30 Hz, coherence was computed using 3 Slepian tapers (time bandwidth product of 2). For frequencies less than 30 Hz, coherence was computed using a single Slepian taper (time bandwidth product of 1). Coherence estimates obtained without the initial step of bandpass filtering data showed similar results.

To compute the correlation between attentional differences in pulvinar spike-cortical field coherence and attentional differences in cortical firing rate, we used the RFs of the pulvinar neurons. These RFs overlapped with the RFs of the cortical neurons.

### Conditional spectral Granger causality analysis

We bandpass-filtered (3-100 Hz) the LFP from each brain area, downsampled to 200 Hz, subtracted the mean, then divided by the standard deviation. For each recording session, we derived a multivariate autoregressive model for each attention condition (attention at the response field for LFPs corresponds to the location of the cue evoking the peak response; attention away from the response field corresponds to the location most far away in the opposite visual hemifield). The autoregressive equation is given by 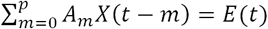, where *A*_*m*_ are the coefficient matrices, *m* is the lag, *X(t)* is the multidimensional process defined for a segment of the time series, and *E(t)* is the noise vector. The model order, *p*, generally corresponded to the first minimum Akaike information criterion value. We used a model order of 10. To estimate *A*_*m*_ and *V*, the covariance matrix of the noise vector, we used the Levinson, Wiggins, Robinson algorithm. To check autoregressive models, we tested the assumption of white model residuals, the stability of the model (i.e., stationary and convergent), and the consistency between the recorded and model-generated data (20). The spectral matrix of the time series is given by *S*(*f*)=*H*(*f*)*VH*\*(*f*),where 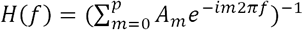 is the transfer function, and ^*\**^ denotes the matrix transpose and complex conjugate.

We calculated conditional Granger causality (21, 22) as a measure of the influence one brain area (Y) has on another area (X), after taking into account additional areas (Z). The conditional Granger causality can be expressed as a function of frequency, to investigate the oscillatory nature of LFPs. In the frequency domain, the conditional Granger causality is given by 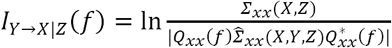, where Σ_xx_(XY)is the variance of the noise in the joint regression of X and Z (variance associated with X), and Q_xx_ and 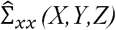 are functions of the transfer function and noise covariance matrix (23, 24). For each attention condition (attention at the response field, defined as the cue location evoking the peak response across areas, and attention away from the response field, defined as the location most far away in the opposite visual hemifield), we calculated the conditional spectral Granger causality in 300 ms sliding windows across the trial. To compare attention conditions across the population, we used *t* tests to determine whether there was significantly greater conditional Granger causality in particular frequency bands (i.e., alpha, beta and gamma) when attention was at the response field location compared to when attention was away from the response field. We controlled the experiment-wise error rate (p < 0.05) using the Holm’s sequential Bonferroni procedure.

---

**Fig. S1.**
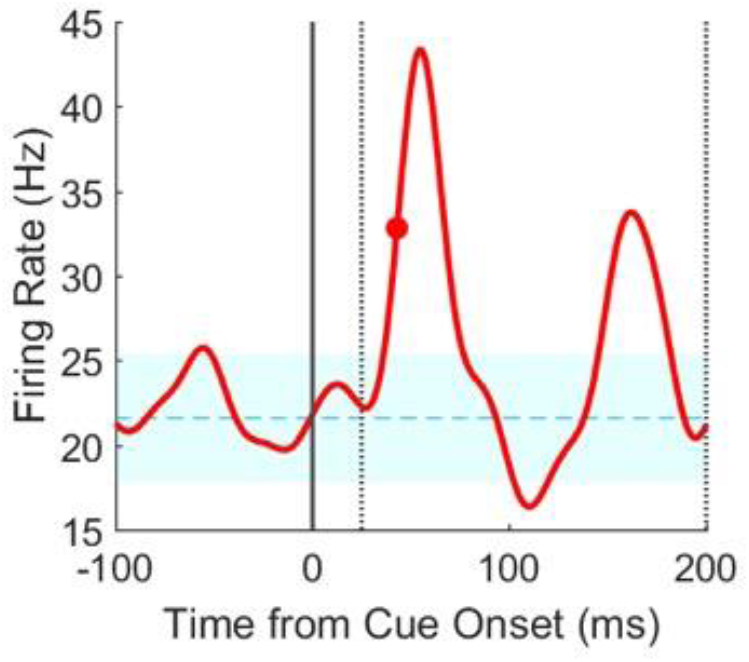
Calculation of response latencies in the pulvinar, V4, and LIP. Latency of firing rate response of an example pulvinar neuron to the cue. Response latency is defined as the time from stimulus onset (cue or array) to when the firing rate reaches half of its extreme rate between 25 and 200 ms after event onset. The extreme rate is the first peak or trough at least 2 standard deviations from the mean baseline rate (dotted cyan line with shading).

**Fig. S2.**
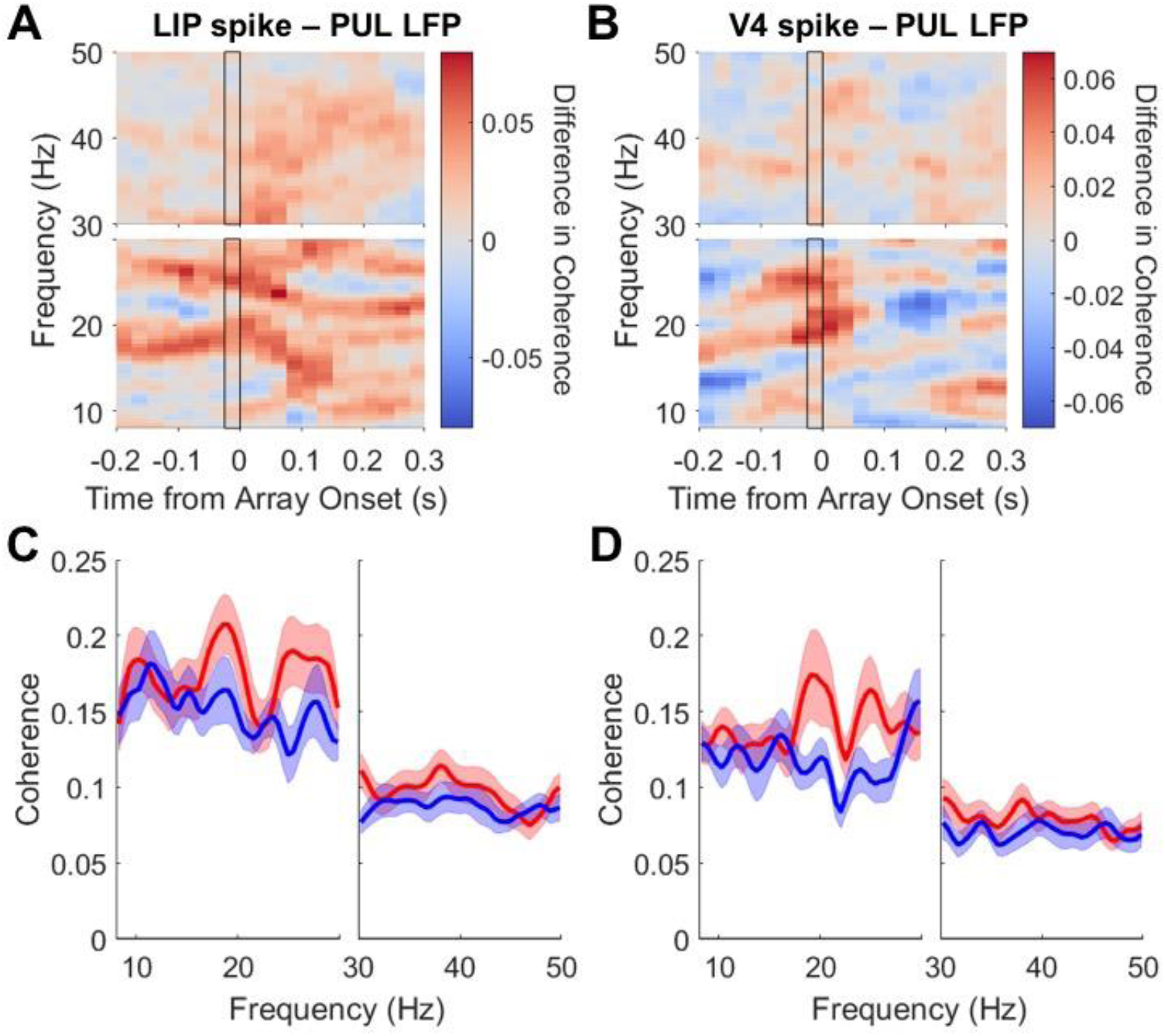
Cortical influence on the pulvinar. **(A-B)** Time-frequency plots of **(A)** LIP spike-pulvinar field coherence, **(B)** V4 spike-pulvinar field coherence. Plots are as in Fig. 4. **(C-D)** Spectra calculated in 0-300 ms window prior to array onset (black-outlined column in A-B). (**C**) LIP spike-pulvinar field coherence. (**D**) V4 spike-pulvinar field coherence.

**Fig. S3.**
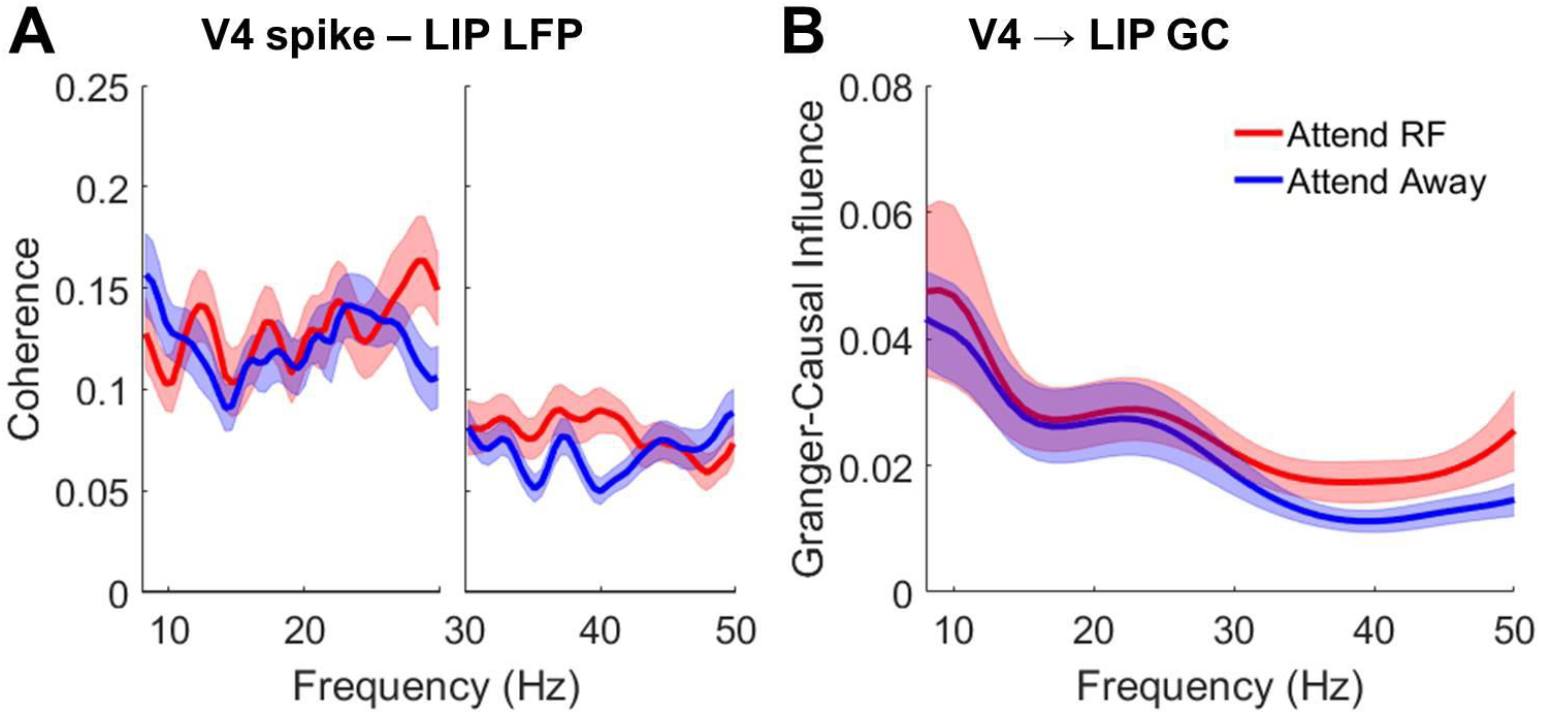
Spatial attention did not modulate V4 influence on LIP. **(A-B)** Spectra calculated in 0-300 ms window prior to array onset. (**A**) V4 spike-LIP field coherence. (**B**) Conditional Granger-causal influence of V4 LFP on LIP LFP (accounting for pulvinar).

